# DNA methylation, microRNA expression profiles and their relationships with transcriptome in grass-fed and grain-fed Angus Cattle rumen tissue

**DOI:** 10.1101/581421

**Authors:** Yaokun Li, José A. Carrillo, Yi Ding, Yanghua He, Chunping Zhao, Jianan Liu, Linsen Zan, Jiuzhou Song

**Affiliations:** College of Animal Science, South China Agricultural University, Guangzhou, Guangdong, P.R. China, 510642; Department of Animal & Avian Sciences, University of Maryland, College Park, MD 20742, USA; College of Animal Science and Technology, Northwest A&F University, Yangling, Shaanxi, P.R. China, 712100

**Keywords:** Rumen, miRNA, methylation, gene expression, grass-fed cattle, grain-fed cattle

## Abstract

Rumen is a critical organ for supplying nutrients for the growth and production of bovine, which might function differently under grass-fed and grain-fed regimens considering the association of gene expression, DNA methylation, and microRNA expression. The objective of this study was to explore the potential mechanism influencing rumen function of grass-fed and grain-fed animals. Methylated DNA binding domain sequencing (MBD-Seq) and microRNA-Seq were respectively utilized to detect the DNA methylation and microRNA expression in rumen tissue of grass-fed and grain-fed Angus cattle. Integration analysis revealed that the expression of the differentially expressed genes ADAMTS3 and ENPP3 was correlated with the methylation abundance of the corresponding DMRs inside these two genes, and these two genes were reported to be respectively involved in biosynthesis and regulation of glycosyltransferase activity; the differentially expressed microRNA bta-mir-122 was predicted to possibly target the differentially expressed genes OCLN and RBM47, potentially affecting the rumen function; the microRNA bta-mir-655 was exclusively detected in grain-fed group; its targets were involved in the significantly enriched insulin and TGF-beta signaling pathways, which might worked together to regulate the function of rumen, resulting in different characteristics between grass-fed and grain-fed cattle. Collectively, our results provided insights into understanding the mechanisms determining rumen function and unraveled the biological basis underlying the economic traits to improve the productivity of animals.

## Introduction

Historically, most of the beef products were from grass-finished cattle. Since the 1950’s, numerous studies were performed to improve the efficiency of beef production; meanwhile, the beef cattle feedlot industry began to emerge, where high-energy grains were utilized to improve the productivity of cattle. And, consumers have been accustomed to grain-fed beef, considering the flavor and palatability resulted from the diets with large proportion of high-energy grain^1^. However, due to new research on the effects of diverse feeding regimens, consumers’ preferences for beef quality have changed, making certain producers to regress to the pastoral beef production in spite of the feeding inefficiency. Recently, growing number of consumers are interested in products obtained from grass-finished animals, raising concerns about the quality difference between grass and grain-fed beef. During the past several decades, studies demonstrated that different feeding regimens could cause alterations in the nutritional composition of beef. Omega-3 and omega-6 are two essential fatty acids, which cannot be synthesized by the body and must be obtained from food; and they are critical for animal health^2^. Studies illustrated that grass-fed beef had a significantly higher content of omega-3 and displayed a more favorable omega-3: omega-6 ratio compared with grain-fed beef^3, 4^. Beta-carotene, the precursor of vitamin A, was more abundant in the muscle of pasture-fed animals than grain-fed ones, which could modulate immune reaction and protect individuals against bacterial and viral infection^5–8^. Meanwhile, grass-finished beef also had higher concentration of diterpenoids and derivatives of chlorophyll, changing the flavor and aroma of the cooked beef^9^. Additionally, higher level of vitamin E was found in the grass-fed beef than the beef products from concentrate diets^10^. Besides nutritional components, the difference regarding the genetics of cattle could also be detected due to diverse feeding regimens^11, 12^.

In the field of genetics, epigenetics refers to cellular and physiological phenotypic trait variations due to external or environmental factors that can switch genes on and off without changing the DNA sequence^13^. Three epigenetic mechanisms, including DNA methylation, histone modifications and small noncoding RNAs, were used to regulate gene expression. Disturbing any of these interacting systems could cause abnormal expression of certain genes. It has been extensively accepted that both DNA methylation at the gene’s promoter region and microRNA (miRNA) regulation are significant in regulating gene expression^14^. Additionally, DNA methylation has been largely studied due to its involvement in the regulation of most biological processes, including embryonic development, genetic imprinting, transcription, chromatin structure, and chromosome stability^15–19^. In mammals, DNA methylation can interfere with transcription factor binding, indirectly repressing the gene activity via recruitment of methyl-CpG binding domain (MBD) protein that can alter chromatin structure^20^. The strength of the repression could depend on the concentration of DNA methylation. In porcine, differentially methylated regions in the promoter have been found to be associated with the repression of both known obesity-related genes and novel genes^21^. Until now, genome-wide DNA methylation profiles of many organisms, including chicken, pig, arabidopsis, human and bovine^21–26^, have been reported. However, the DNA methylation pattern of bovine rumen remains less studied.

Additionally, miRNAs could also regulate gene expression at the post-transcriptional level. They specifically bind to corresponding mRNAs, leading to RNA silencing and translation repression. It is known that each miRNA might have many mRNA targets, and each mRNA could also be regulated by more than one miRNA. In mammals, miRNAs are predicted to regulated the expression of ∼60% of all protein-coding genes, and found to be associated with many biological processes, including cell cycle control, cell growth and differentiation, apoptosis and embryo development^27, 28^.

As the largest part of the stomach, rumen constitutes the main site for plant material digestion and microbial fermentation. It plays a critical role in supplying nutrition for animals’ maintenance, growth and production. In the present study, we hypothesized that rumen may function differently under grass-fed and grain-fed regimens, which could result in different compositions and flavor of beef. Given the emerging roles of miRNAs and DNA methylation in gene regulation, identifying the DNA methylation profile and expression pattern of miRNAs is critical to understand their functions in the process of bovine development. Therefore, we utilized Illumina sequencing technology to characterize the genome-wide miRNA expression pattern and DNA methylation profile in the rumen tissue of grass-fed and grain-fed Angus cattle. Further, we performed the integrated genome-wide analysis of miRNAs, DNA methylation, and mRNA expression, which was important for future functional study and the discovery of valuable epigenetic biomarkers. The findings of this study would be significant for identifying potential mechanisms that led to differences observed between grass-fed and grain-fed cattle.

## Materials and Methods

### Ethics Statement

All animal experiments were conducted according to the NIH guidelines for housing and care of laboratory animals and in accordance with the regulations of the University of Maryland at College Park (UMCP). The UMCP Institutional Animal Care and Use Committee (IACUC) reviewed and approved the protocols (permit number R-08-62).

### Sample collection

The studied Angus cattle were born and raised at the Wye Angus farm, which has been closed for almost 75 years and yielded genetically similar progenies. It provides an excellent experimental resource to perform biological research. The genetic similarity permits us to better control the cause of variation among experimental individuals. We randomly chose pairs of animals from larger sets of steers that received a particular treatment. All animals in the study received the same diet until weaning. Then, we randomly assigned the male calves to one diet and exclusively received that regimen until termination. The grain-fed group received conventional diet comprised of shelled corn, corn silage, soybean and trace minerals. The grass-fed steers normally consumed grazed alfalfa; during wintertime, bailage was supplied. The alfalfa was harvested from the land and cultivated without any fertilizers, pesticides or other chemicals. The grass-fed animals ate no animal, agricultural or industrial byproducts and never received any type of grain. The grain–fed animals reached the market weight around the age of 14 months; however, grass-fed steers needed approximately 200 additional days to reach the similar weight value with the age of 20 months. Immediately after termination at the Old Line Custom Meat Company (Baltimore, MD), the rumen samples from all experimental animals were excised at the same location around the cardiac ostium, and then the samples were rinsed and preserved at −80°C for subsequent analysis.

### DNA extraction and MBD-Seq library construction

Firstly, we extracted the genomic DNA from two samples of each group using the Wizard Genomic DNA purification kit (Promega, A1120). The DNA concentration was measured by the Qubit dsDNA Broad-Range Assay (Invitrogen, Q32850), which was then adjusted to 0.1μg/μl for a final volume of 55 μl in a 1.5 ml tube and sheared into 300–500 bp fragments using the Bioruptor Sonicator; the fragmented DNA was checked on agarose gel to visualize the size of the resultant segments. Secondly, MethylCap kit (Diagenode, C02020010) was employed to obtain the methylated DNA. The 141.8 μl of capture reaction mix containing 12 μl of sheared DNA was prepared according to the instruction of the kit. Then, 119 μl of capture reaction mix was incubated with 1 μl of diluted MethylCap protein at 40 rpm on a rotating wheel for 2 hours at 4°C to let the interaction occur. The rest (22.8 μl) was used as input sample. Next, meDNA Capture beads (coated with GSH) provided by the kit were utilized to capture methylated DNA. Unbound DNA was washed off and the bead pellet combined by the methylated DNA was collected. After washing the bead pellet, we performed the elution of the captured DNA; 150 μl of low, medium, and high concentration elution buffer were sequentially used per capture reaction. All fractions and input were purified using the MiniElute PCR Purification Kit (QIAGEN, 28006).

NEBNext End Repair Module (NEB, E6050S, USA) was used for the end repair of the fragmented methylated DNA. Then the 3’ poly “A” was added through DNA Polymerase I, Large (Klenow) Fragment (NEB, M0210L, USA). Next, a pair of Solexa adaptors (Illumina) was ligated to the repaired ends by T4 ligase (Promega, M1801, USA). The ligated products were electrophoresed in 2% agarose gels; the fragments (DNA plus adaptors) from 200 to 500bp were selected and purified by QIA- quick Gel Extraction Kit (QIAGEN, USA). Then, we enriched the purified DNA templates through PCR using Phusion Hot Start High-Fidelity DNA Polymerase (NEB, M0530S, USA). After purification of the PCR products (MinElute PCR Purification Kit, QIAGEN, USA), the concentration of the DNA library was measured through the Qubit assay (Life Technology, Q32850). Finally, we performed DNA sequencing in the Solexa 1G Genome Analyzer (Illumina) following the specification provided by the manufacturer.

### MBD-Seq data analysis

The quality of the raw reads was firstly evaluated employing FastQC, which is a web-based software that thoroughly examines the reads and creates a detailed and extensive quality assurance report containing the indicators including “per base sequence quality”, “per base sequence content”, and “per sequence GC content”, etc. Then, Bowtie aligned the reads to the reference genome (Bos_taurus_UMD3.1/bosTau6), which was downloaded from the iGenomes web site (http://support.illumina.com/sequencing/sequencing_software/igenome.html). During this step, according to the indicators generated by FastQC, we trimmed the first 10 bases and the last 5 bases of each read (50 bp) to maintain high sequence quality score, which resulted in 35 bp tags. For data format conversion, SAMtools and BEDtools were applied in our analysis.

For peaks identification, the Model Based Analysis of ChIP-Seq (MACS) was implemented individually for each sample. MACS models the shift size of tags and improves the spatial resolution of predicted binding sites. This software also utilizes a dynamic Poisson distribution to effectively catch the local bias, improving the reliability of the prediction. Identification of the differentially methylated regions (DMR) was performed by the R package Diffbind; it calculates the differentially bound sites using affinity data. The input data for DiffBind was the bam file containing the aligned reads and the peaks set identified by MACS. For normalization, the default method TMM (Trimmed Mean of M-values) considering the effective library size was used. After creating a contrast between conditions, DiffBind carried out a DESeq2 analysis with a false discovery rate (FDR) < 0.1 to call the DMRs. DESeq2 approach employed shrinkage estimation for fold changes and dispersions, making the estimated values more stable and interpretable^29^.

The ChIPpeakAnno package was used for the genomic annotation of the previously identified DMRs^30^. This software provides information about the distance, relative position and overlaps for the inquired feature. The annotation information was obtained from BioMart, using Ensembl 80 in the archive site; dataset “Bos taurus_genes_ensembl (UMD3.1)” corresponding to the bosTau6 reference genome was used for alignment. The CpG islands annotation was retrieved from the UCSC web browser. Finally, the DMRs were annotated based on specific genomic features. In addition, we integrated the results of DMRs and transcriptome which has been published in 2015^31^, and compared the methylation abundance of the DMRs and the expression level of their corresponding genes, from which we hypothesized the relationship between the methylation abundance of the DMRs and the expression level of their corresponding genes.

### Bisulfite Sequencing for MBD-Seq Validation

After quality evaluation and quantification, equal amounts of DNA from two samples of each group were pooled together, serving as the template for the bisulfite conversion and the bisulfite PCR. Firstly, sodium bisulfite conversion reagents were used to treat 500 ng of each DNA pool (Methyl EdgeTM Bisulfite Conversion System, Promega, USA). The DMRs for validation were randomly selected from the bioinformatics analysis results. The PCR primers were designed through MethPrimer (http://www.urogene.org/cgi-bin/methprimer/methprimer.cgi) and shown in Supplemental Table 1. Then, PCR products were purified using QIA- quick Gel Extraction Kit (QIAGEN, USA), which were subsequently ligated to pGEM-T Vector (pGEM-T Vector System I, Promega, USA), and transformed to DH5α competent cells (Z-Competent E. Coli Cells—Strain Zymo 5α, ZYMO Research, USA) for screening successful insertions (blue-white selection) after incubation at 37°C overnight. Next, ten white colonies from each culture plate were cultured overnight in 37°C shaker. Plasmid DNA was isolated using Zyppy Plasmid Miniprep Kit (ZYMO Research, USA). BigDye Terminator v3.1 Cycle Sequencing Kit (Applied Biosystems, 4337456) was employed for sequencing in the ABI 3730 machine. Bisulfite sequencing results were analyzed by QUMA (http://quma.cdb.riken.jp). Finally, DNA methylation level for each region and group was obtained.

### RNA extraction and microRNA-Seq library construction

Total RNA was extracted individually (three animals per group) using Trizol (Invitrogen, Carlsbad, CA, USA) followed by DNase digestion and Qiagen RNeasy column purification (Invitrogen, Carlsbad, CA, USA), as previously described^32^. The RNA sample was dissolved in RNAse-free H2O; then we checked the integrity and quality by a NanoDrop 1000 spectrophotometer and by visualization on a 1.5% agarose gel. The microRNA-Seq library was constructed using NEBNext Multiplex Small RNA Library Prep Set for Illumina (Set 1) following the instructions provided by the manufacturer (NEB, E7300S/L, USA). The library construction was started with 800ng total RNA. Firstly, the 3’ adaptor and 5’ adaptor were sequentially ligated to the RNA. Then, we performed the reverse transcription to synthesize the first strand. Next, we executed the PCR amplification and added the 6-bp index to the DNA products; different libraries were assigned different indexes. Subsequently, we selected and recycled the DNA fragments from 140 to 150 bp using 6% polyacrylamide gel (6% Novex® TBE PAGE gel, Life Technology, USA), which were corresponding to RNAs from 21 to 30 bp. After purification, the concentration of the library was measured through the Qubit assay (Life Technology, Q32850). Finally, the libraries identified by the 6-bp index were sequenced at 50 bp/sequence read using an Illumina HiSeq 2000 sequencer, as described previously^33^.

### microRNA-Seq data analysis

After quality evaluation of the tags performed by FastQC, all samples were analyzed employing miRDeep* software, which could quantify known and novel miRNAs from small RNA sequencing. This program had a user-friendly graphic interface implemented totally in java and accepted raw data in FastQ and SAM/BAM format as input. The default length of miRNAs was set at 18–23 nucleotides. Low-quality reads were filtered out at the alignment stage; the read with more than a 20 phred score was considered as good read. Meanwhile, multi-mapping reads with alignments to more than 100 genomic loci were also filtered out; the score based on the probabilistic score of the potential miRNA precursor was set at −10 as default^34^. In the further miRNA expression analysis, we only considered known miRNAs. Expression values for known miRNAs were estimated individually by miRDeep* in each sample. Subsequently, these expression levels were extracted and recorded in a matrix, which was used later as input for the edgeR package to call the differentially expressed miRNAs.

After identification of the differentially expressed miRNAs, the target genes for those miRNAs were predicted using TargetScan, which were then utilized for the integrated analysis with transcriptome analysis results published in 2015^31^.

### Quantitative real-time PCR (qRT-PCR) analysis

qRT-PCR was conducted to validate the differentially expressed miRNA found in the microRNA-Seq analysis on the iCycler iQ PCR system (Bio-Rad, Hercules, CA, USA). The RT-PCR reactions were performed with a QuantiTect SYBR Green PCR Kit (Qiagen, Valencia, CA, USA) according to the manufacturer’s instructions. Three technical replicates and two independent biological replicates were performed for each product. RPS18 was selected as the control gene. The primer sequences were listed in Supplemental Table 2.

## Results

### Landscape of the DNA methylomes

For the methylome study, we had four experimental samples in total; the alignment levels were 94.92%, 82.98%, 95.54% and 96.10%, respectively (Supplemental Fig. 1). We finally found 217 DMRs between the two groups (Supplemental Table 3), which were presented as red dots in Fig. 1. Based on the identified DMRs, the cluster analysis of our experimental samples was performed (Fig. 2), the results showed consistency with the group assignment of the samples, suggesting that these DNA fragments carrying DMRs could be used to predict the potential mechanism causing the difference between our two groups. From those DMRs, we found that only two DMRs were highly methylated and all the others low methylated in the grain-fed bovines compared with the grass-fed group. For the distribution of the DMR length, the average was 3760 bp with extreme value of 485–bp and 14,387 bp. Approximately one percent of the DMRs were less than 1,000 bp and 37.8% of the DMRs accounted for fragments longer than 4,000 bp (Fig. 3). Fig. 4 showed the number of DMRs per chromosome, chromosome 8 accounted for 18 DMRs indicating the largest number, following by chromosome 1 with 15 DMRs. Chromosomes 19, 21, 23, and 15 had the lowest number of DMRs with values of 2, 2, 1 and 1, respectively.

**Figure 1.**
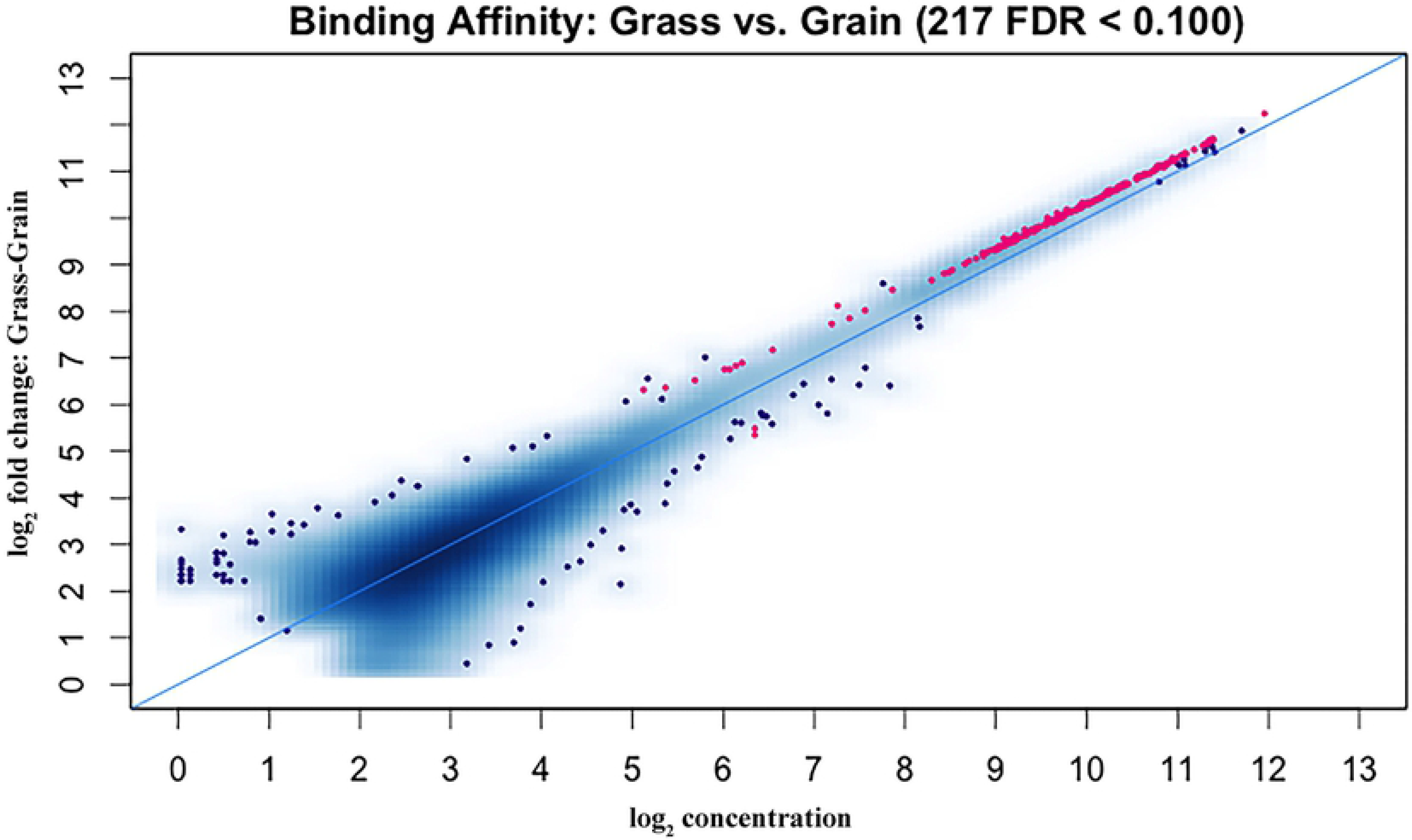
DMRs between grass-fed and grain-fed steers. The MA plot shows in red the DMRs obtained with a false discovery rate of < 0.1.

**Figure 2.**
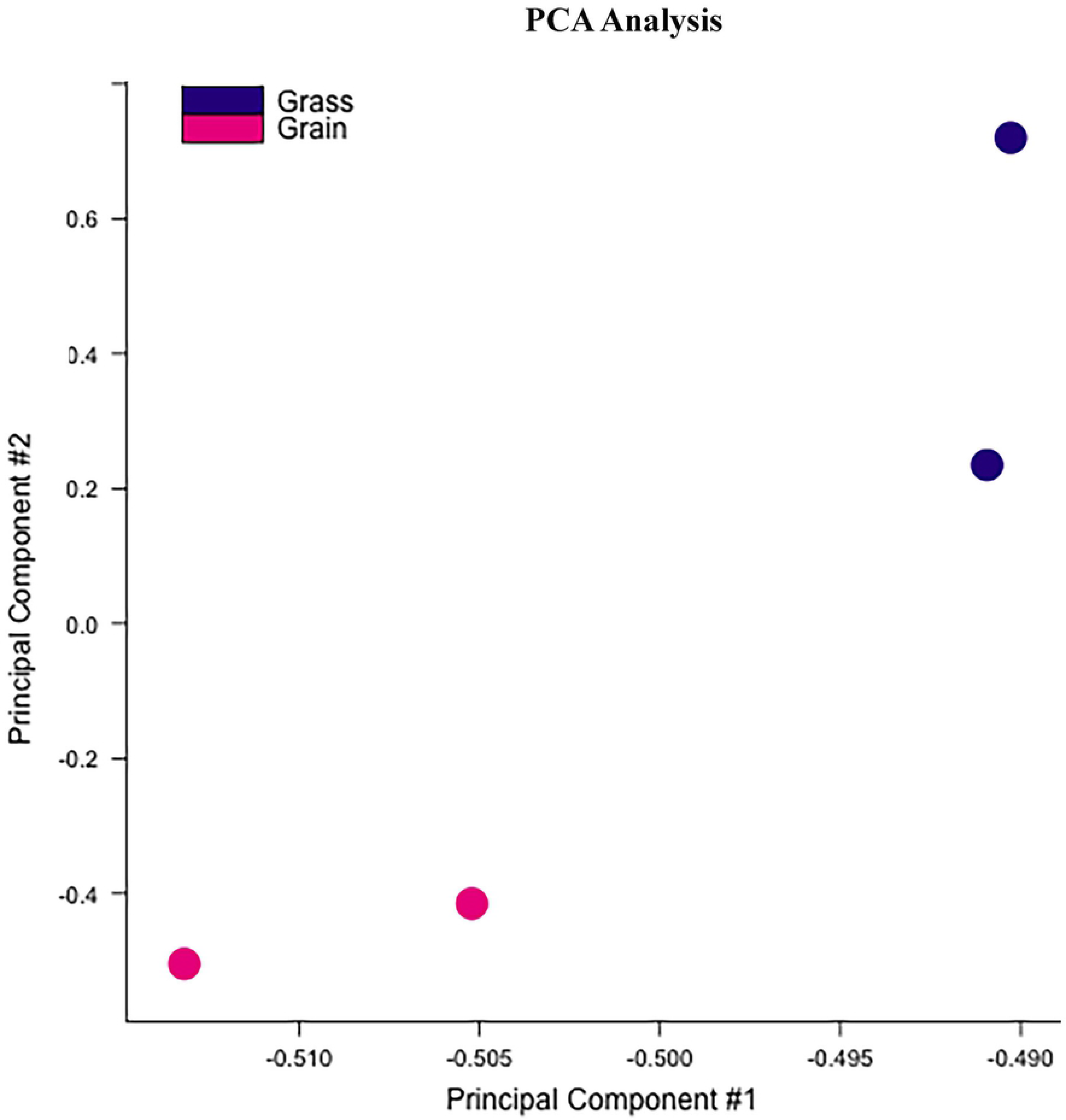
PCA analysis of the experimental individuals based on the identified DMRs. PCA: principle component analysis.

**Figure 3.**
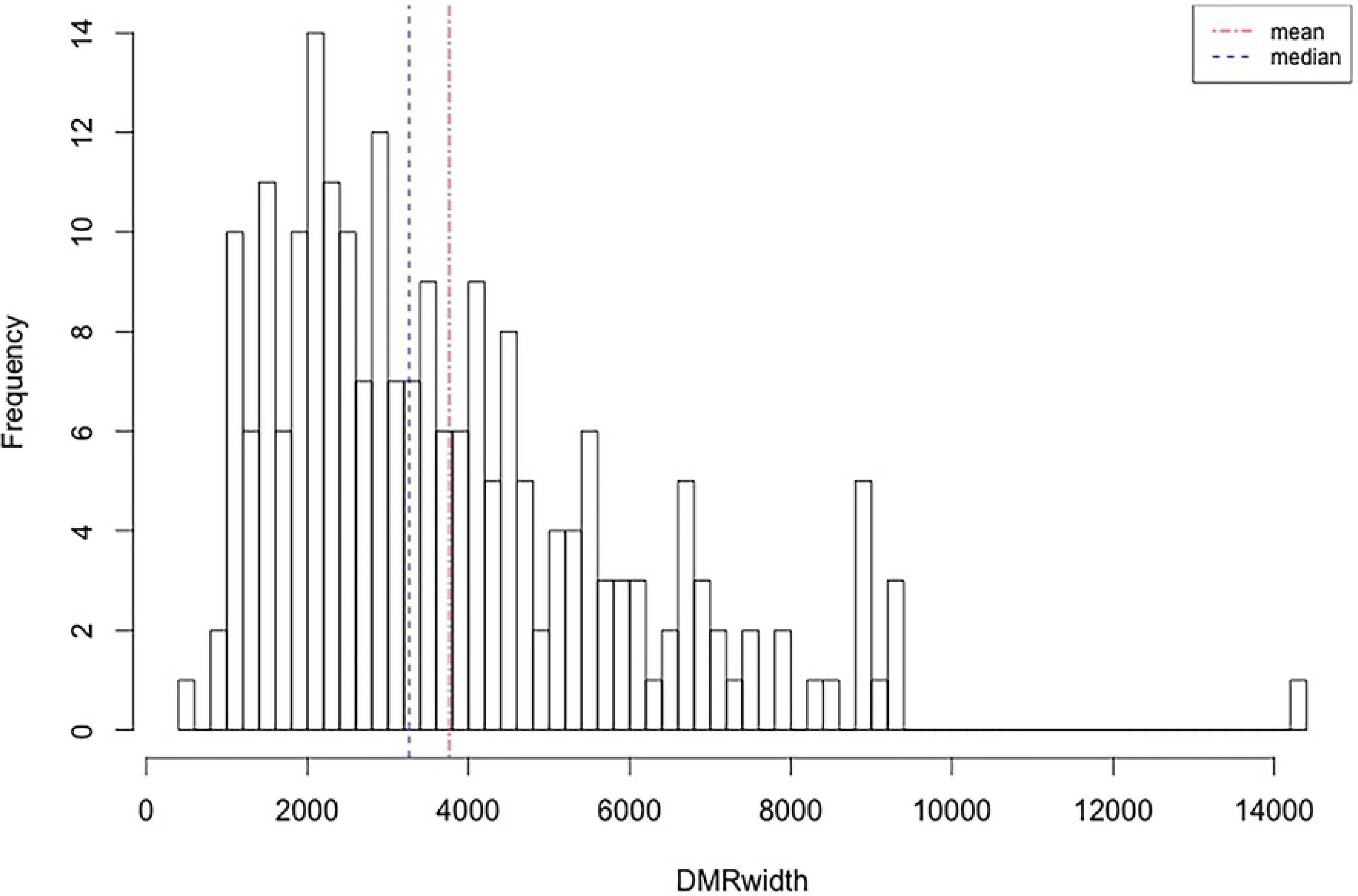
DMRs’ length density. The abscissa represents the extent of the DMRs in base pairs. The dotted light blue and red lines correspond for the median and mean respectively.

**Figure 4.**
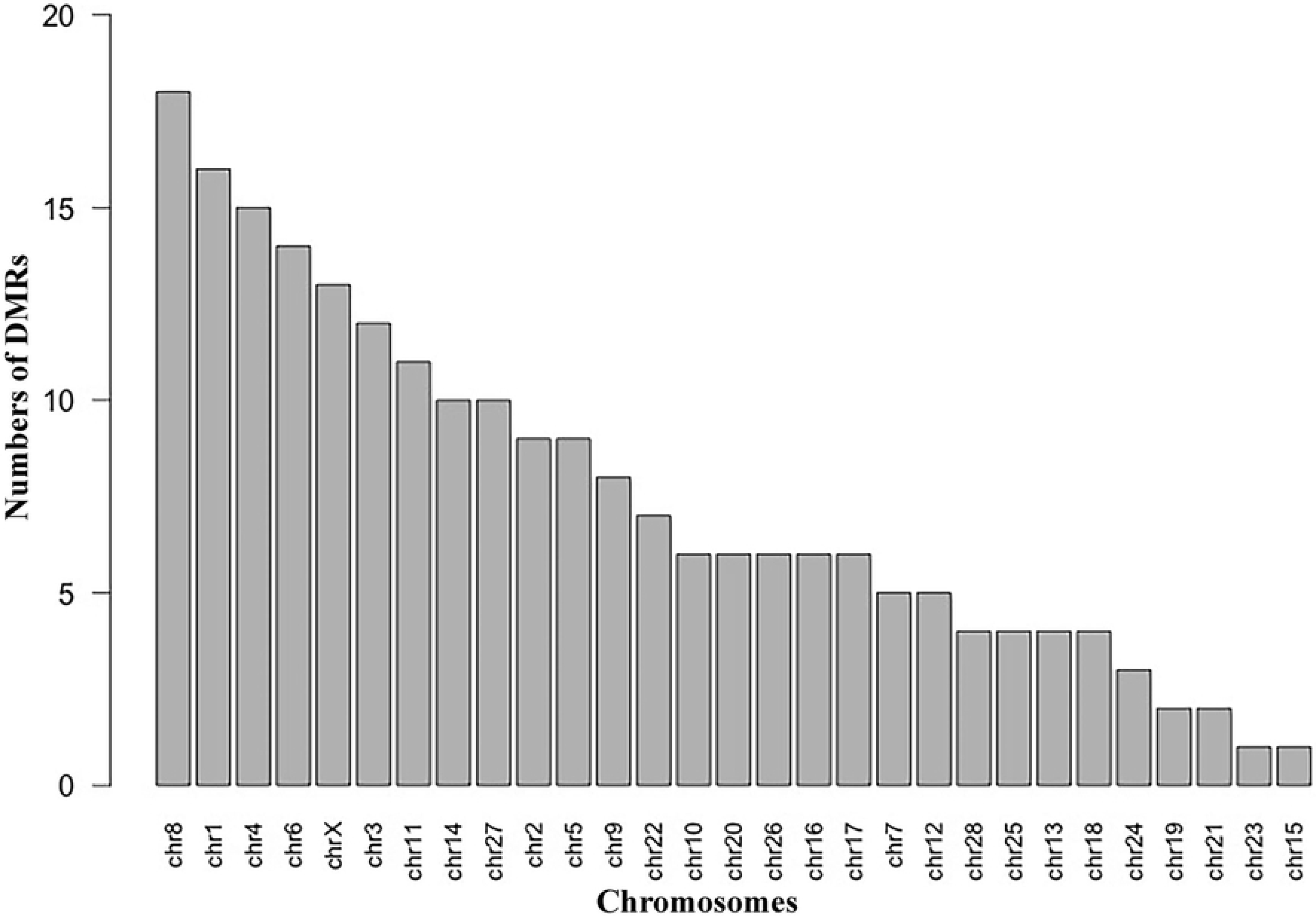
Chromosomal location frequencies of the DMRs. Distribution of the DMRs per chromosome without normalization (ignoring chromosome length).

The binding affinity between grass-fed and grain-fed cattle was shown in Fig. 5. We found that global DNA methylation level decreased in grain-fed individuals. Then, we detected the DNA methylation level in the 2 kb region upstream of the transcription start site (TSS), the 2 kb region downstream of the transcription end site (TSS) and the gene body region from TSS to TES. In our experimental groups, the DNA methylation level declined significantly before the TSS and increased notably towards the gene body region with slight changes before the TES, followed by a sharp decrease in the downstream of TES (Fig. 6). Compared with grass-fed individuals, the grain-fed steers showed a higher level of DNA methylation around the gene body region.

**Figure 5.**
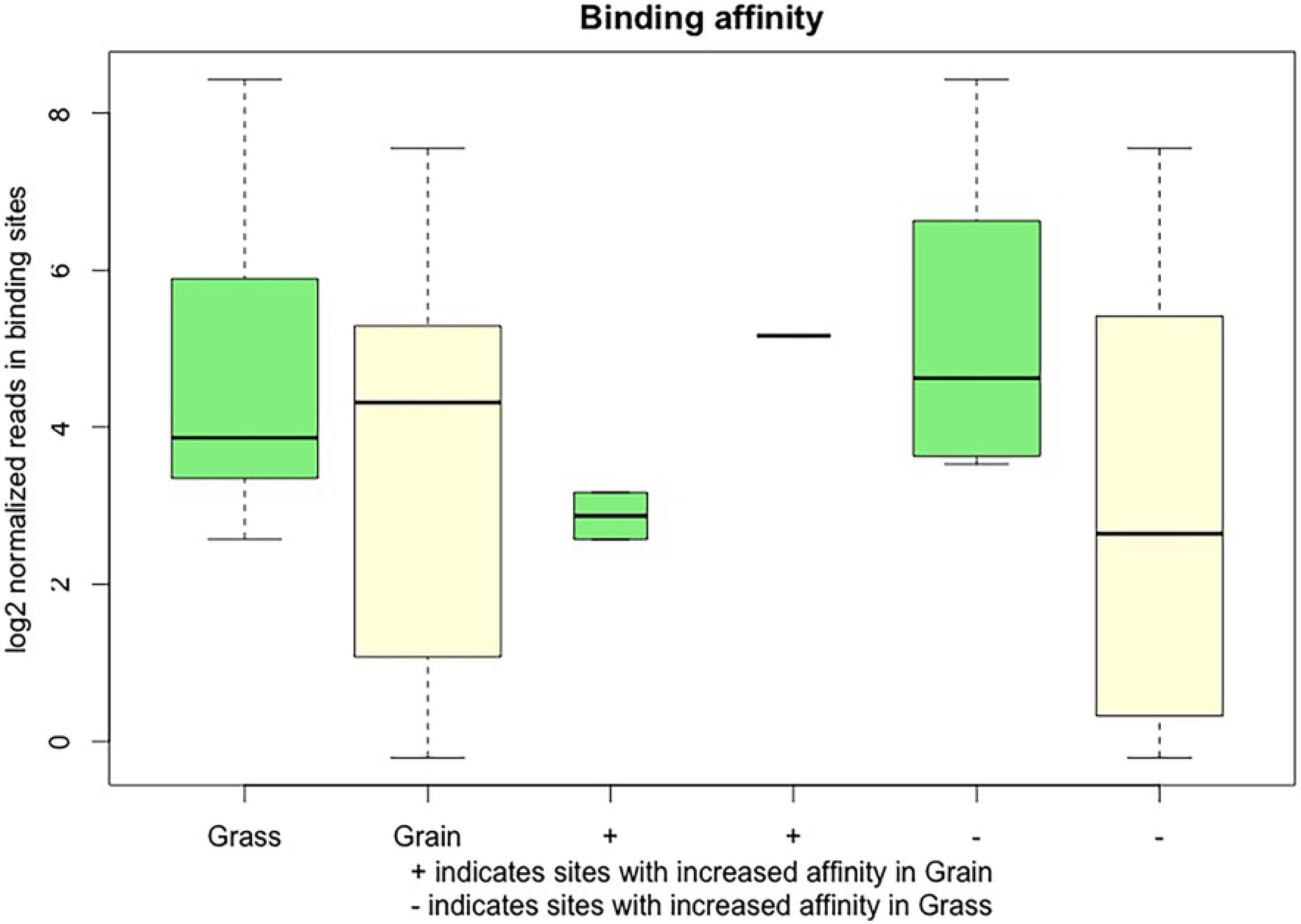
Normalized reads at the binding sites per condition. The first two boxplots represent the overall methylation level in the grass-fed and grain-fed groups. The + sign marked the sites with increased binding affinity in grass-fed group, the − sign represented the regions that enhanced the binding ability in the grain-fed group. The light green and yellow boxes corresponded to grass-fed and grain-fed with + and −, respectively.

**Figure 6.**
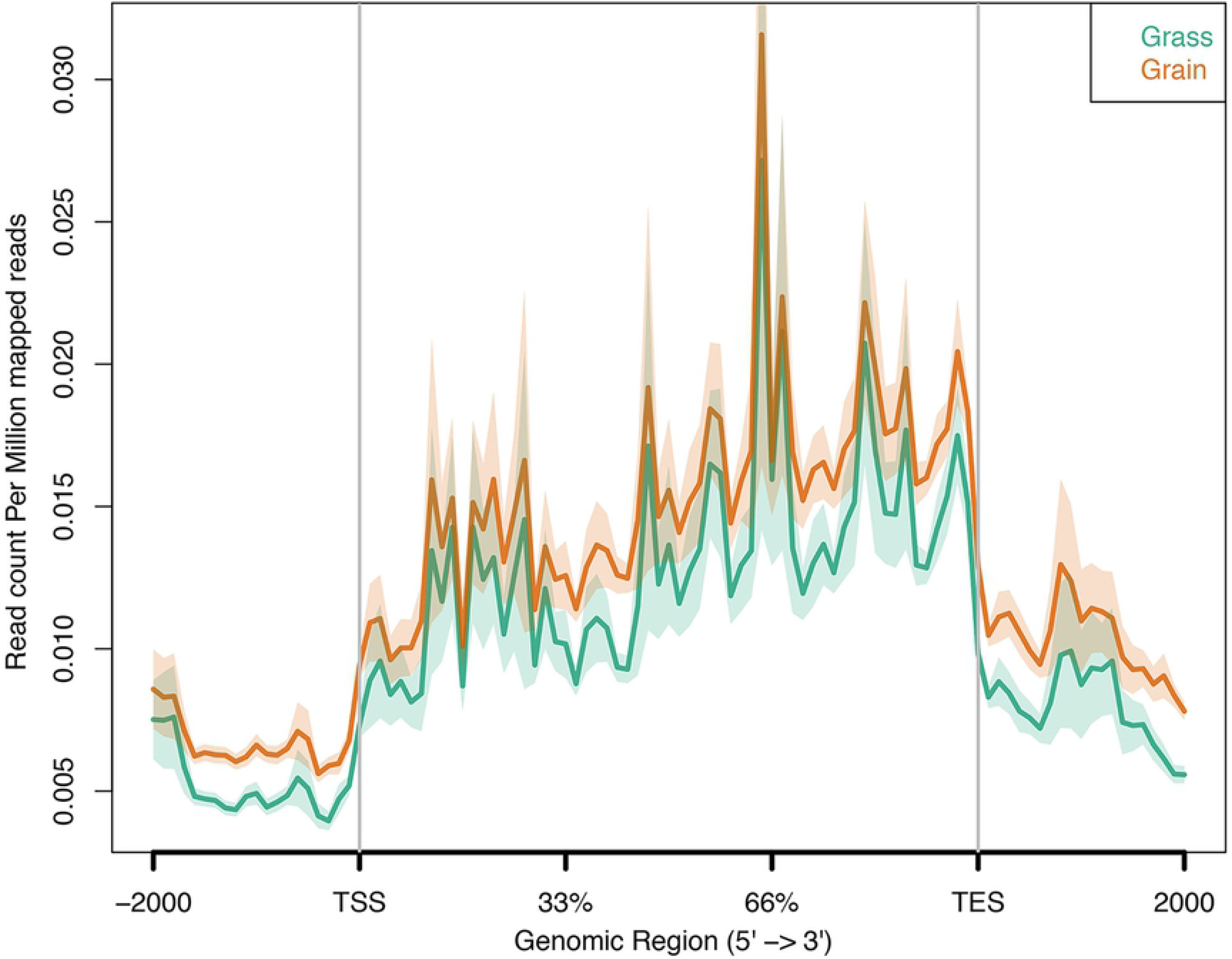
DNA methylation patterns around gene bodies in grass-fed and grain-fed animals measured by MBD-Seq. The x axis indicates the position around gene bodies, and the y axis indicates the normalized read number aligned to the normalized gene body region and the around region.

The DMR annotation was subsequently performed (Supplemental Table 4). The distribution of DMRs’ distance to the nearest TSS could be found in Fig. 7, suggesting that approximately 14.3% of the DMRs are located within a range of 10,000 bp from the TSS. Moreover, DMRs’ location related to genes and CpG islands were summarized in Fig. 8. We found that almost a quarter of the DMRs were contained within genes and half in regions upstream of the TSS. The DMRs downstream of the genes accounted for 25%. Only four genes were found in the DMRs. Regarding CpG islands, approximately 8.8% of the annotated CpG islands were included in the DMRs. Most of the DMRs were outside of the CpG island boundaries with 42.4% upstream and 50% downstream of the CpG islands.

**Figure 7.**
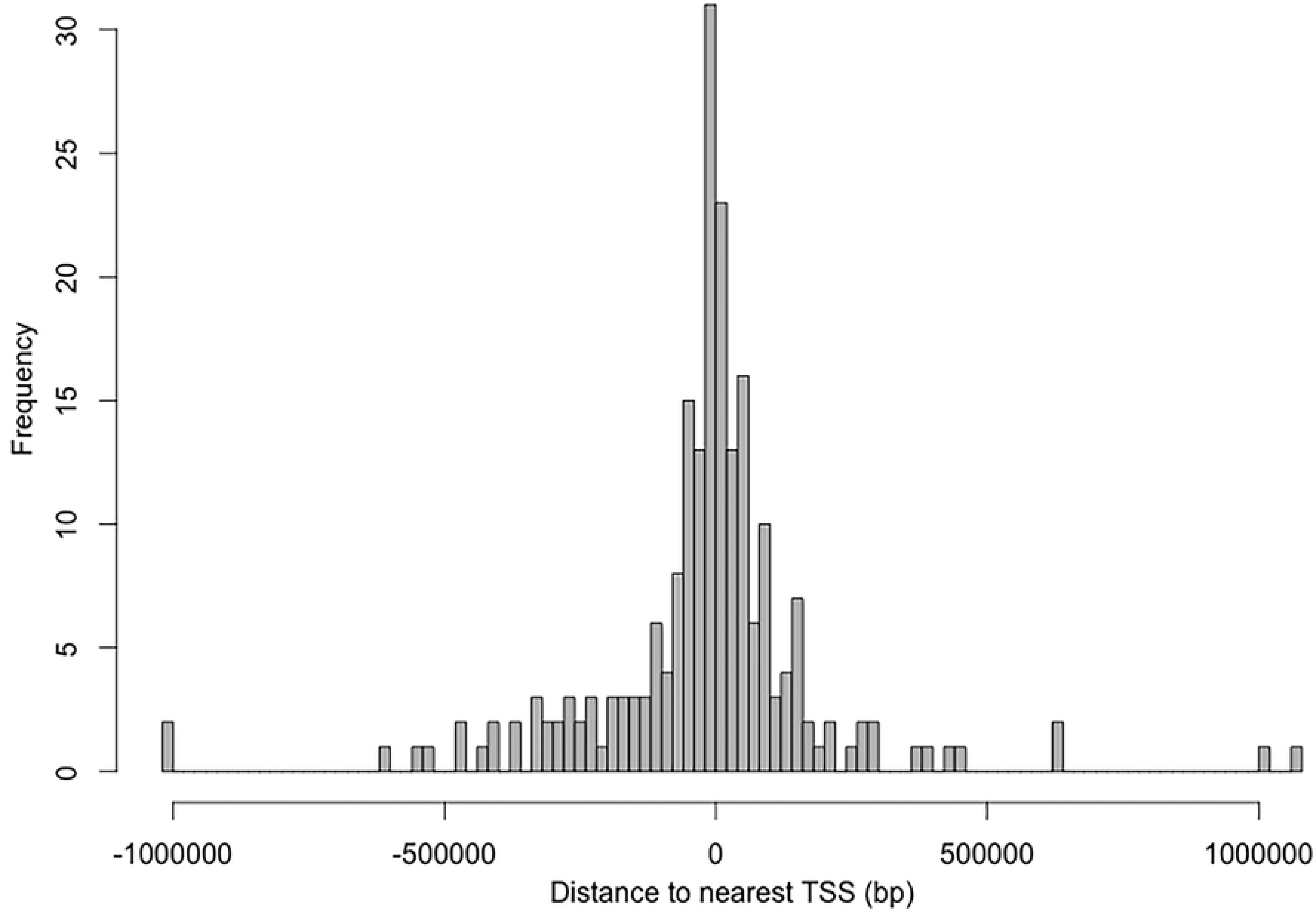
Frequency of DMRs’ distance to closest transcription start site. The distance from the transcription start site is represented in base pairs from the 0 in the x axis.

**Figure 8.**
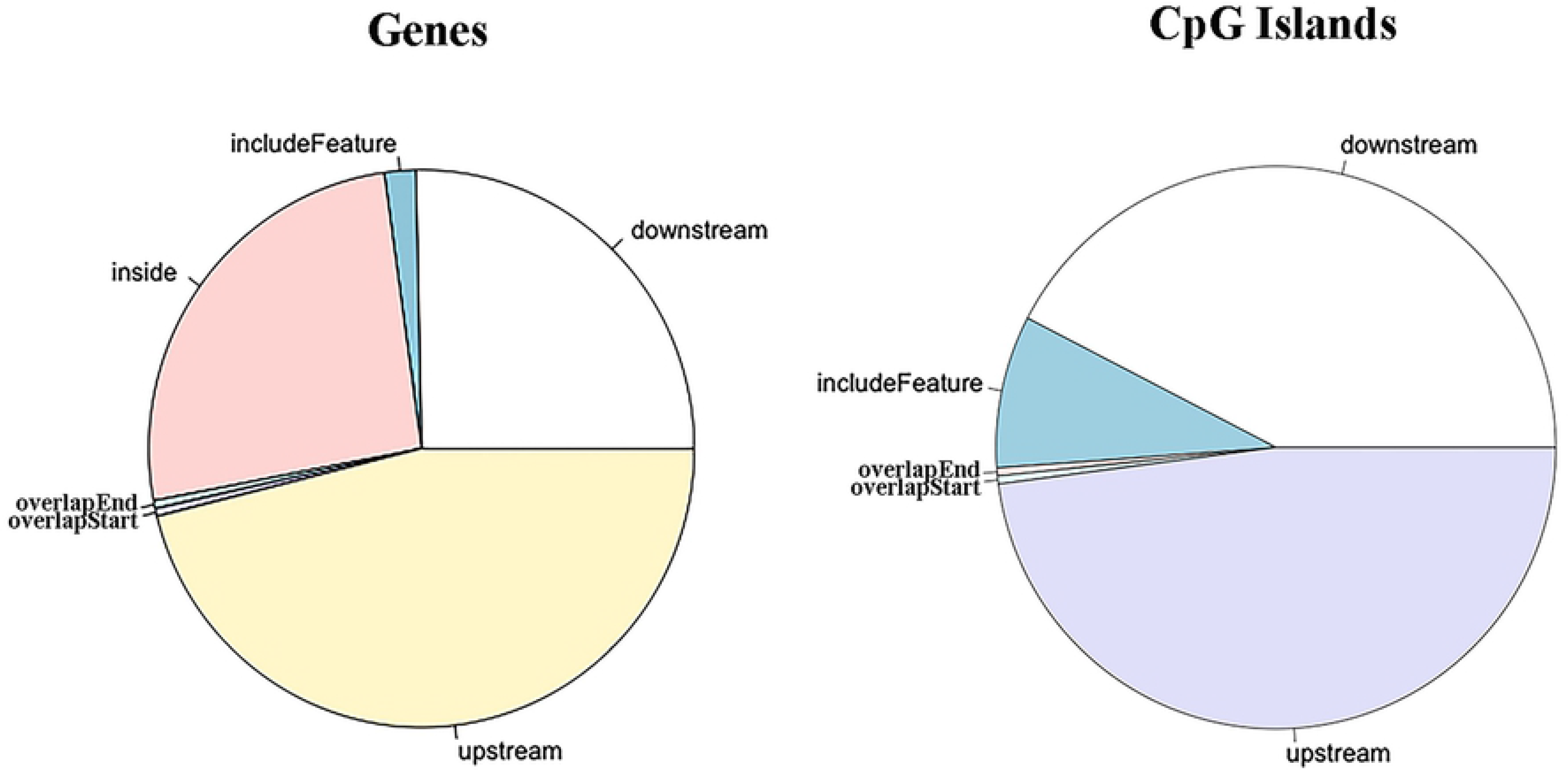
DMRs location distribution regarding genes and CpG islands. The labels in both pies represent: inside, DMR contained within the feature; include feature, the genomic feature is entirely included in the DMR; overlap start, the DMR extend over the start site of the feature; overlap end, the ending site of the feature overlaps with the DMR; downstream, DMR locates downstream the feature; upstream, the DMR aligns upstream the genomic feature.

### Integrated analysis of MBD-Seq with transcriptome (RNA-Seq)

In this study, we screened the genes with promoters containing DMRs, using 10,000 bp as the parameter for maximum distance. From the 217 identified DMRs, 21 were located within the promoters, corresponding to 21 different genes. Our previous study explored the transcriptome profiling in grass-fed and grain-fed animals, and 342 differentially expressed genes (DEGs) were detected between the two groups^33^. We found that, of the 21 genes, only the expression of 8 genes was positive correlated with the corresponding DMRs, the other 13 genes were negative correlated with the DMR, however, none of the 21 genes could be found in the DEGs list. Additionally, 57 DMRs were found inside 52 genes, and all of the DMRs were highly methylated in grass-fed steers. And two of the corresponding genes were detected in the DEGs list; one was negative correlated with the corresponding DMR, and the other one was positive correlated with the DMR, which were ADAMTS3 and ENPP3, respectively. Generally, we found that 47.3% of the corresponding 52 genes were positive correlated with the DMR. Meanwhile, we found 4 genes were inside the DMRs; however, none of them were differentially expressed. Based on the annotation regarding the TSS, another two genes could also be discovered in the DEGs list, which were CRISPLD1 and PRR5; CRISPLD1 was negative correlated with the corresponding DMR and PRR5 was positive correlated with the DMR; the nearest distance of the corresponding DMR to the TSS was respectively 14,973 and 45,887 bp.

### MBD-Seq and microRNA-Seq data validation

In order to assess the reliability and accuracy of DMRs detection from MBD-Seq, 5 regions were randomly selected for validation. The calculated methylation levels are shown in the Fig. 9. We found that the methylation level of all the 5 regions agreed with MBD-Seq analysis. The bisulfite sequencing employed only a segment of the methylation region to perform the validation, thus the magnitude of methylation difference of the validation results could not be exactly the same as MBD-Seq analysis.

**Figure 9.**
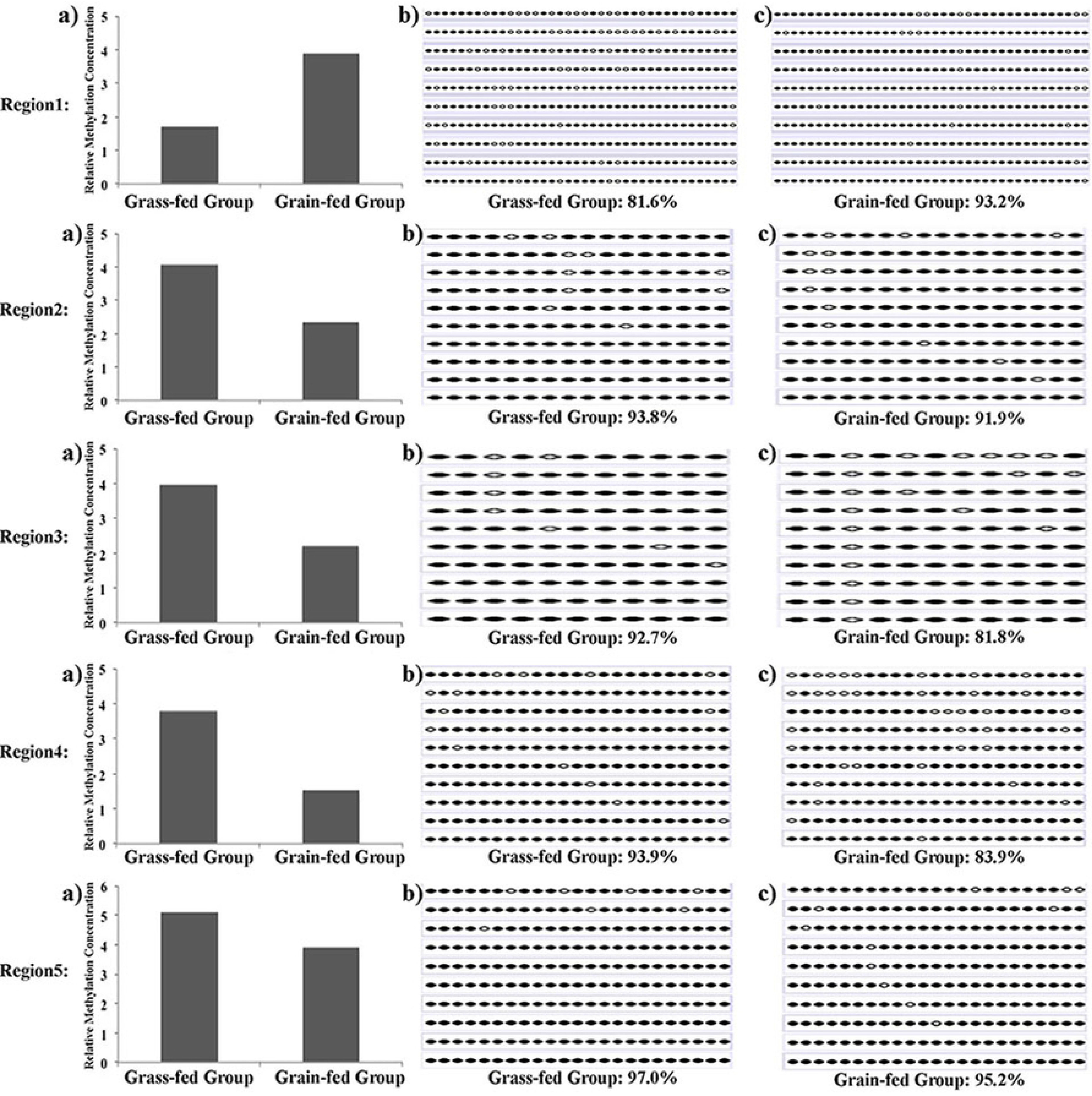
Bisulfite sequencing validation of MBD-Seq results. a) Methylation concentration levels from MBD-seq. b) and c) Bisulfite sequencing results. Each line represents a plasmid sequence and each dot indicates a CpG site. An open circle indicates an unmethylated CpG site and a black dot stands for one methylated CpG site. The methylation level was calculated as the number of methylated CpG sites divided by the total detected CpGs.

For the differentially expressed miRNA bta-mir-122, the expression level in grain-fed ruminal wall was significantly higher than in grass-fed ruminal wall, and the results of qPCR and RNA-Seq suggested the same direction (Supplemental Fig. 2).

### Expression profiles of microRNAs in grass-fed and grain-fed groups

To identify the microRNAs involved in bovine rumen function, total RNAs from three biological replicates for each condition were used to construct small RNA libraries. Via high-throughput sequencing, we obtained an average of 15 million reads per microRNA sample. The percentage of alignment for all samples exceeded 90% as shown in Figure 11. Based on the aligned sequencing reads, we totally identified 321 known miRNAs (Supplemental Table 5), of which 72.9% were high expressed in grain-fed individuals compared with grass-feed ones. Between the two groups, we identified only one differentially expressed miRNA with FDR value less than 0.1, which was bta-mir-122 with improved expression in grain-fed group as compared with grass-fed animas. Additionally, we found that microRNA bta-mir-655 was exclusively expressed in grain-fed group.

### Target gene prediction, Gene Ontology and pathways enriched by the differentially expressed microRNAs

Prediction of target genes of the miRNA was performed using TargetScan (http://www.targetscan.org/). For bta-mir-122 and bta-mir-655, we respectively found 145 and 749 target genes. To explore the specific functional features shared by the targets of the above miRNA, online software David Bioinformatics Resources 6.7 was used to perform the GO enrichment analysis, mainly regarding the biological processes, cellular components and molecular functions. For the targets of bta-mir-122, the most significant GO terms were: regulation of GTPase activity, regulation of protein signal transduction, intracellular, GTPase activator activity and GTPase regulator activity (Table 1). To some extent, the targets were also involved in the pathway glycolysis/gluconeogenesis (P = 7.9×10^−2^), which could produce energy and is an critical metabolic pathway employed by a host of organisms. Therefore, this might be an interesting pathway to study in rumen. For the targets of bta-mir-655, the most significant GO terms in the three categories were: regulation of cellular process, biological regulation, developmental process, regulation of metabolic process, intracellular and transcription factor activity (Table 2). The complete list of GO terms enriched with the target genes of bta-mir-655 was provided in Supplemental Table 6. Then, we detected the pathways that involved the targets of bta-mir-655, the most significantly enriched pathways were shown in Table 3. Several interesting pathways were found including adherens junction, insulin signaling pathway, TGF-beta signaling pathway and neurotrophin signaling. These results would provide prior knowledge to explain the difference between grass-fed and grain-feed Angus cattle.

**Table 1.**
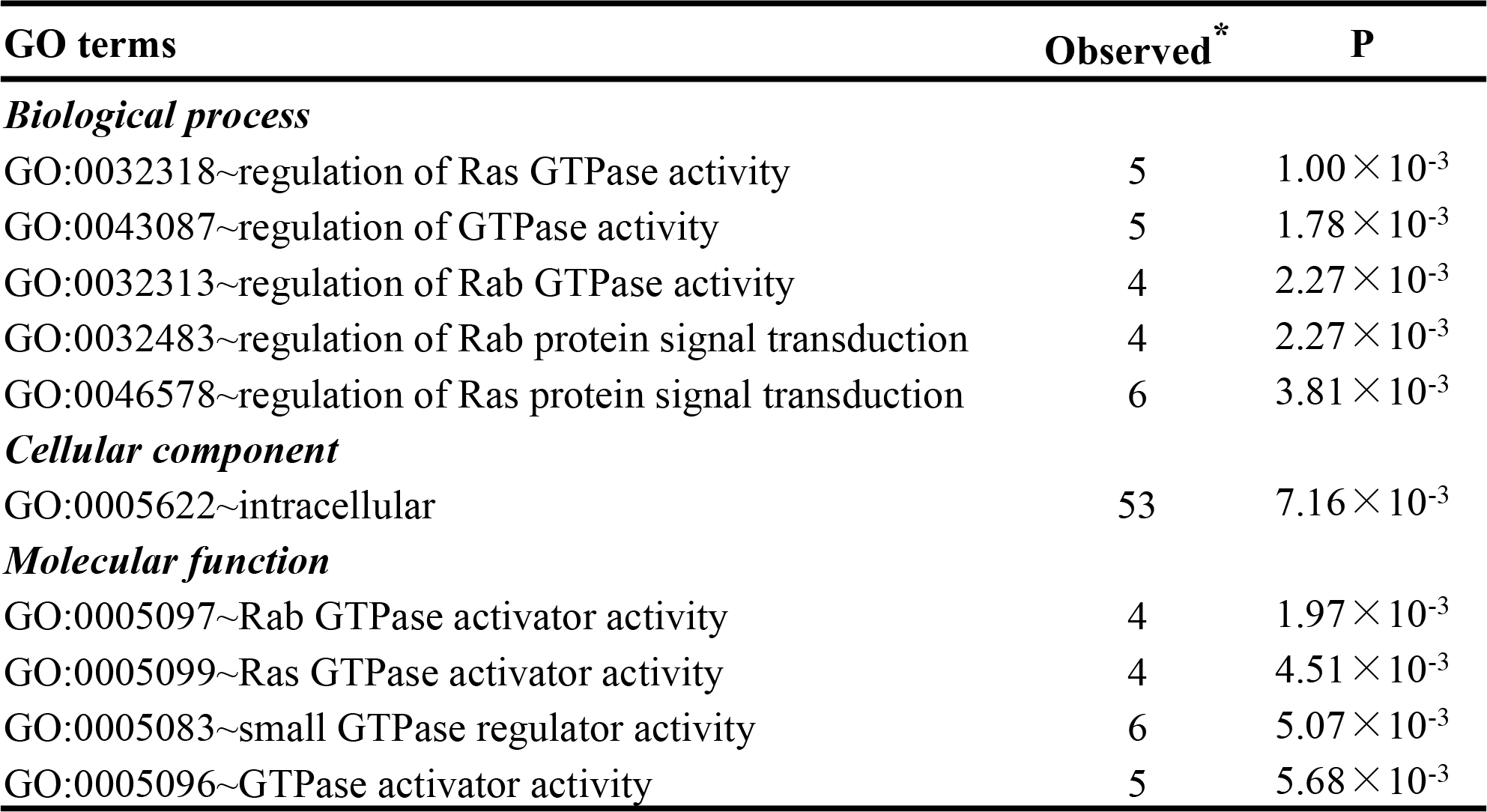
Gene Ontology (GO) terms enriched with the targets of bta-mir-122 (P < 0.01)

| GO terms | Observed* | P |
| --- | --- | --- |
| ***Biological process*** |  |  |
| GO:0032318~regulation of Ras GTPase activity | 5 | $1.00 \times 10^{-3}$ |
| GO:0043087~regulation of GTPase activity | 5 | $1.78 \times 10^{-3}$ |
| GO:0032313~regulation of Rab GTPase activity | 4 | $2.27 \times 10^{-3}$ |
| GO:0032483~regulation of Rab protein signal transduction | 4 | $2.27 \times 10^{-3}$ |
| GO:0046578~regulation of Ras protein signal transduction | 6 | $3.81 \times 10^{-3}$ |
| ***Cellular component*** |  |  |
| GO:0005622~intracellular | 53 | $7.16 \times 10^{-3}$ |
| ***Molecular function*** |  |  |
| GO:0005097~Rab GTPase activator activity | 4 | $1.97 \times 10^{-3}$ |
| GO:0005099~Ras GTPase activator activity | 4 | $4.51 \times 10^{-3}$ |
| GO:0005083~small GTPase regulator activity | 6 | $5.07 \times 10^{-3}$ |
| GO:0005096~GTPase activator activity | 5 | $5.68 \times 10^{-3}$ |

**Table 2.**
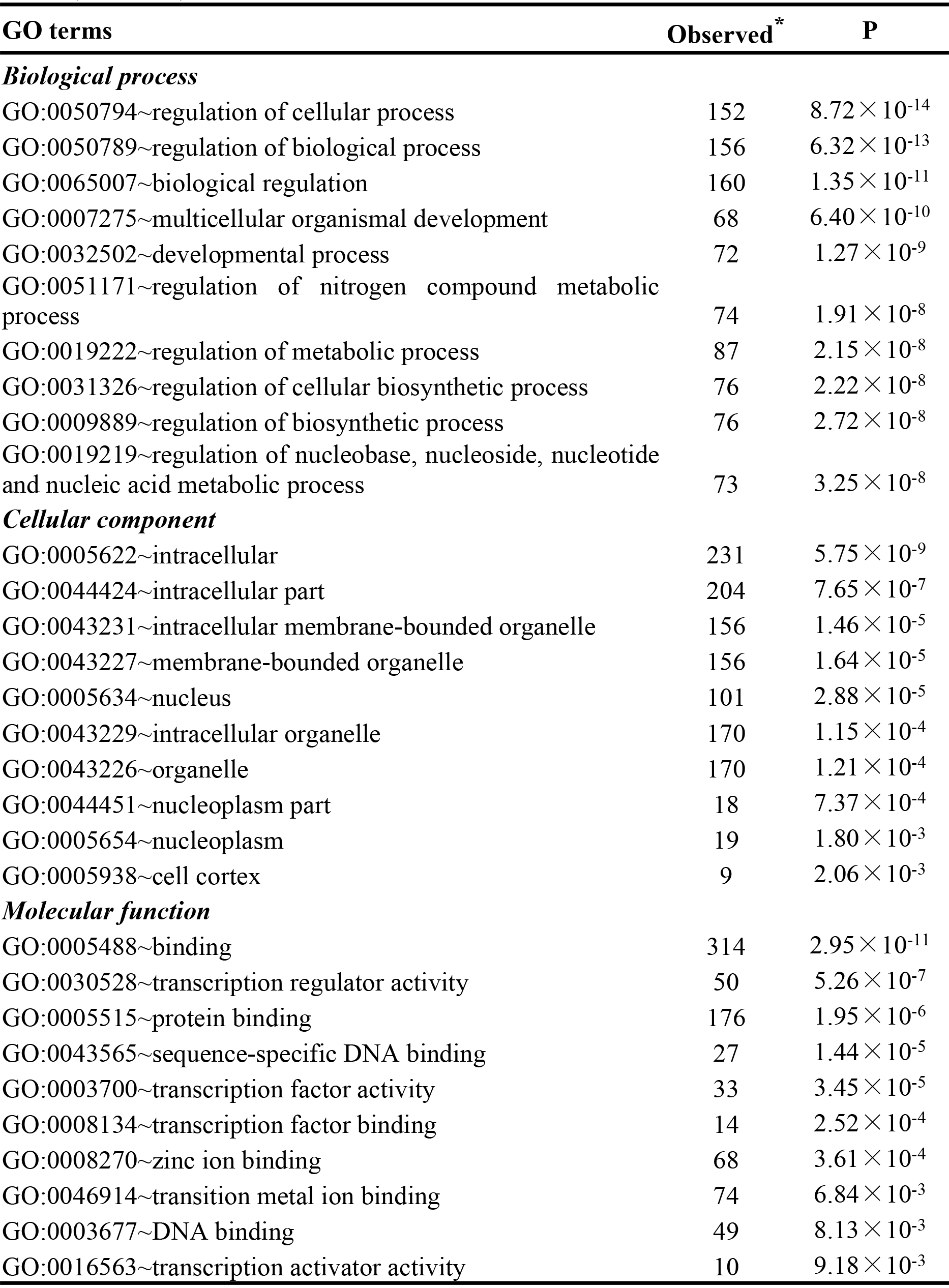
The top Gene Ontology (GO) terms enriched with the targets of bta-mir-655 (P < 0.01)

**Table 3.** KEGG pathways enriched with the target genes of bta-mir-655 by GO analysis (P < 0.01)

| KEGG Pathway | Observed* | P value |
| --- | --- | --- |
| Adherens junction | 10 | $2.32 \times 10^{-4}$ |
| Insulin signaling pathway | 13 | $5.49 \times 10^{-4}$ |
| TGF-beta signaling pathway | 9 | $3.35 \times 10^{-3}$ |
| Neurotrophin signaling pathway | 11 | $4.03 \times 10^{-3}$ |
| Renal cell carcinoma | 8 | $4.24 \times 10^{-3}$ |
| Pancreatic cancer | 8 | $5.03 \times 10^{-3}$ |
| Fc gamma R-mediated phagocytosis | 9 | $5.64 \times 10^{-3}$ |
| Pathways in cancer | 19 | $7.07 \times 10^{-3}$ |
| Axon guidance | 10 | $9.87 \times 10^{-3}$ |

\*Number of the target genes in the pathway

### Integrated analysis of microRNA-Seq with transcriptome (RNA-Seq)

For bta-mir-122, two of the 145 target genes were found in the DEGs list from RNA-Seq, which were OCLN and RBM47, respectively. These two genes were all highly expressed in grass-fed group, showing a negative correlation with bta-mir-122. For bta-mir-655 exclusively expressed in grain-fed animals, we found that 14 of the 749 targets were overlapped with the DEGs. Among the 14 overlapped genes, DPF1, GAS1, ANO1 and NFASC displayed increased expression in grain-fed group, suggesting positive correlation with bta-mir-655; the other nine genes (MYO10, SASH1, EMP1, SLC14A1, PCDH19, IRX5, MAL2, FAM84A, AHCYL2 and DSG3) exhibited enhanced expression level in grass-fed group, which demonstrated negative correlation with bta-mir-655.

## Discussion

In addition to genetic makeup, age, species, gender and environment, grass and grain ratio of the diet could also contribute to significant difference in the general profile of fatty acid, animals’ growth rate, the content of vitamin A and E in the meat, the antioxidant enzyme concentration in body tissues and lipid depots^35–37^. In the past several years, miRNA and DNA methylation have been studied widely. However, limited studies have focused on the rumen tissue of bovine, one of the most significant livestock in the world. Rumen was the most important workshop for the digestion of nutritional substance of bovine. The potential changes of rumen metabolism may have effects on the quality and quantity of protein, affecting other digestive organs, such as reticulum, small intestine and large intestine. Therefore, rumen function was critical for animals’ growth, health and productivity. In the present study, we mainly focused on the rumen under different feeding diets, detecting the genome-wide DNA methylation profiles and miRNA expression profiles, and we also performed integrated analysis of these with transcriptomic results. The objective was to identify the DMRs and miRNAs that might potentially affect the rumen function. This study performed a comprehensive analysis of DNA methylation profiles in the rumen tissues of grass-fed and grain-fed animals, and totally revealed 217 DMRs across the whole genome. In our analysis, we found that 21 DMRs were located in the promoter regions; however, none of their corresponding genes were overlapped with the DEGs. For these DMRs, although the methylation level was different, the concentration might not be high enough to cause the expression difference of their targets. Or, these DMRs could actually exert no effects on the genes. Additionally, we detected 57 DMRs inside 52 genes, of which two genes (ADAMTS3 and ENPP3) were overlapped with the DEGs. The methylation abundance of the DMRs exhibited negative and positive correlation with the expression of the corresponding gene ADAMTS3 and ENPP3, respectively. Studies suggested that DNA methylation in the gene body region might change the chromatin structure and alter the transcription elongation efficiency^18^. Gene body methylation was more prevalent than promoter; its relationship with gene expression levels was very complex and not monotonic. Normally, gene body DNA methylation showed positive correlation with gene expression^38^. However, the information about the role of DNA methylation in the gene body region was still insufficient. For the two DEGs predicted to be correlated with the DMR in the gene body region, ADAMTS3 belonged to the ADAMTS (a disintegrin and metalloproteinase domain with thrombospondin motifs) metalloproteinase family mediating cartilage aggrecan degradation as well as collagen biosynthesis^39^. It was involved in the biosynthesis of type II procollagen, the main collagen of articular cartilage^40^. ENPP3 was known to be a typical ectoenzyme localized to the cell surface, playing a role in metabolizing extracellular nucleotides and their derivatives^41^. And, it was suggested that ENPP3 could modulate nucleotide-mediated signal transduction, which was known as purinergic signaling^42, 43^. Recent study demonstrated that ENPP3 could catalyze the hydrolysis of the nucleotide sugar, and the levels of several intracellular sugars, including UDP-Fuc, UDP-GalNAc and UDP-GlcA, were significantly influenced by knocking down endogenous ENPP3^44^. There was evidence showing that intracellular sugars played a role in regulating glycosyltransferase activity and controlling the total cellular glycosylation profile^45–47^. Therefore, through DNA methylation regulation, the expression of ENPP3 would be changed, which might indirectly regulate the activity of a broad range of glycosyltransferase and consequently influence the total cellular glycosylation pattern.

The miRNAs could negatively regulate gene expression through degrading the target mRNAs or inhibiting the translation. However, the understanding of the contribution of miRNA to rumen function was still limited. In this study, we aimed to get a deeper insight into underlying mechanism of miRNA in rumen function. Between grass-fed and grain-fed group, we found one differentially expressed miRNA (bta-mir-122) with higher expression level in grain-fed steers. Bta-mir-122 was a conserved miRNA between vertebrate species. It was also found differentially expressed between normal and aberrant placental samples, suggesting that bta-mir-122 might be involved in the development of placentae^48^. In liver, bta-mir-122 was relevant in maintenance of homeostasis and had critical metabolic and anti-inflammatory functions^49^. In our study, we detected 145 target genes for bta-mir-122 in total. To explore the specific functional features shared by the 145 targets, we performed GO enrichment analysis. Results showed that the targets were mainly involved in the GTPase activity and regulation of protein signal transduction. GTP hydrolysis played an essential role in controlling numerous biological processes, including protein biosynthesis, growth control and differentiation, and various transport processes. Additionally, two of the 145 target genes were found in the DEGs list, which were OCLN and RBM47, respectively. OCLN was a key tight junction protein that could interact with intracellular signaling pathways, which played roles in regulating intestinal function^50^. RBM47 was a novel RNA-binding protein; it could contribute to the basic machinery for C to U RNA editing in intestine, influencing the expression of some genes^51^; however, the function of this gene has not been studied in rumen. We made the hypothesis that RBM47 would alter the expression of some genes in rumen, consequently regulating rumen function. Meanwhile, we detected one exclusively expressed miRNA (bta-mir-655) in grain-fed group; it could target 749 different genes. After the GO analysis based on the targets, the most significant GO terms included regulation of cellular process, biological regulation, developmental process, regulation of metabolic process, and transcription factor activity, which played a critical role in rumen function. The most interesting pathways were found including adherens junction, insulin signaling pathway and TGF-beta signaling pathway, which were related to animals’ growth, survival and development. In addition, 14 of the 749 targets were overlapped with the DEGs; however, none of them was discovered in the pathways. Thus, we hypothesized that bta-mir-655 just slightly changed the expression of the targets without significant expression change; but the target genes working together in the corresponding pathway could exert significant effects on the rumen function. Among the 14 overlapped genes, FAM84A, localized in the subcellular membrane region, was involved in invasion and/or metastasis of colon cancer cells influencing colorectal cancer^52^; SASH1 was a member of the SH3-domain containing expressed in lymphocytes (SLY1) gene family, it encodes signal adapter proteins which were composed of certain protein–protein interaction domains, showing prognostic significance in human cancer^53^. A previous study suggested that epithelial membrane protein 1 (EMP1) gene could prevent tumor proliferation and was associated with gastric carcinoma^54^. DSG3 was predicted to be involved in the GO term cell adhesion; it was reported to be associated with oncogenesis^55^. However, information about the function of these genes in rumen was still limited; they might play a potential role in animal development through affecting the gastrointestinal function. Accordingly, it might be of great interest to perform functional experiment of these genes to better understand the mechanisms causing the varied performance.

Our results provided evidence for explaining the molecular mechanisms leading to the differences between grass-fed and grain-fed cattle. For the function of the DEGs containing the DMR and the differentially expressed miRNAs, extensive experimental validation work was still needed. Thus, overexpression and inhibition of our identified DEGs and miRNAs could be considered for the functional validation, which would provide more supportive information for our findings.

## Conclusions

In this study, we integrated DNA methylation and miRNA expression with transcriptome analysis to explore the potential mechanism influencing rumen function of grass-fed and grain-fed animals. We found that the expression of ADAMTS3 and ENPP3 might be altered by the corresponding DMR inside these two genes, through which rumen function of grass-fed and grain-fed cattle could be regulated. For the differentially expressed miRNA bta-mir-122, it might modulate the rumen function by targeting the two DEGs OCLN and RBM47, which were possibly associated with gastrointestinal function. While expanding the scope of future studies with putative genes relevant to bovine growth and meat quality traits, our integrated analysis provided biological insights into the mechanisms regulating rumen function and uncovered the molecular basis underlying the economic traits to enhance the productivity of animals.

## Competing interests

The authors declare that there are no competing interests about this manuscript.

## Funding

The work was supported by China Scholarship Council (CSC), Maryland Agricultural Experiment Station (MAES) and Jorgensen Endowment Funds

## Acknowledgment

We thank MAES and China Scholarship Council for their supports of this study.

## Author contributions

Conceived and designed the experiments: L.S.Z and J.Z.S. Performed the experiments: Y.K.L, J.N.L and Y.H.H. Analyzed the data: J.A.C. Contributed reagents/materials/analysis tools: J.Z.S., C.P.Z and G.E.L. Wrote the paper: Y.K.L., J.A.C. and Y.D.

## Additional information

**Supplemental Figure 1: Alignment level of MBD-Seq reads to the Bovine Genome.**

**Supplemental Figure 2: Validation of differentially expressed miRNA.** The mean value of log2 (fold-change) for each group was compared in the bar chart. FC means fold-change.

**Supplemental Table 1:** Primers used for validating the randomly selected DMRs (XLS).

**Supplemental Table 2:** Primers used for validating the differentially expressed miRNA (XLS).

**Supplemental Table 3:** The DMRs between grass-fed and grain-fed Angus cattle (XLS). The threshold of FDR <0.1 was used to call the significant difference

**Supplemental Table 4:** Annotation of the identified DMRs (XLS).

**Supplemental Table 5:** The known miRNAs shared by grass-fed and grain-fed animals (XLS).

**Supplemental Table 6:** The GO terms enriched with the target genes of bta-mir-655 (XLS).

